# Mammal virus diversity estimates are unstable due to accelerating discovery effort

**DOI:** 10.1101/2021.08.10.455791

**Authors:** Rory Gibb, Gregory F. Albery, Nardus Mollentze, Evan A. Eskew, Liam Brierley, Sadie J. Ryan, Stephanie N. Seifert, Colin J. Carlson

## Abstract

Host-virus association data form the backbone of research into eco-evolutionary drivers of viral diversity and host-level zoonotic risk. However, knowledge of the wildlife virome is inherently constrained by historical discovery effort, and there are concerns that the reliability of ecological inference from host-virus data may be undermined by taxonomic and geographical sampling biases. Here, we evaluate whether current estimates of host-level viral diversity in wild mammals are stable enough to be considered biologically meaningful, by analysing a comprehensive dataset of discovery dates of 6,571 unique mammal host-virus associations between 1930 and 2018. We show that virus discovery rates in mammal hosts are still either constant or accelerating, with little evidence of declines towards viral richness asymptotes in even highly-sampled hosts. Consequently, inference of relative viral richness across host species has been unstable over time, particularly in bats, where intensified surveillance since the early 2000s caused a rapid rearrangement of species’ ranked viral richness. Our results show that comparative inference of host-level virus diversity across mammals is highly sensitive to even short-term changes in sampling effort. We advise caution to avoid overinterpreting patterns in current data, since our findings suggest that an analysis conducted today could feasibly draw quite different conclusions than one conducted only a decade ago.

## Introduction

Pathogens are unevenly associated with hosts across the tree of life, and understanding the coevolutionary processes that drive these patterns is important for both fundamental ecological and health-motivated research. For example, data on how viral diversity is distributed across species and geographies can provide insights into biogeographical trends and anthropogenic drivers of cross-species transmission and disease emergence (1–3). Researchers have developed numerous hypotheses about the mechanisms underlying differences in virus diversity across hosts, from broad macroevolutionary trends (e.g., bats were found to be infected by a greater apparent diversity of viruses than other mammal orders (4)) to narrower ecological associations (e.g., longer-lived bats living in larger groups host a greater apparent diversity of viruses (5)). Such work frequently investigates the number of viruses known to infect a given host species (i.e. viral richness), based on synthetic datasets of known host-virus associations that are often compiled for this kind of hypothesis testing (4, 6, 7).

However, recent work has raised concerns that such datasets inspire false confidence in analytical inferences. Although host-virus association datasets take an increasingly complete inventory of current scientific knowledge (8), a substantial proportion of known viruses remain excluded due to long lead times before official taxonomic recognition, which itself is not uniform across the virome (9). An even greater proportion of the global virome remains completely undescribed (10, 11), with current knowledge strongly influenced by discovery strategies (12). There are concerns that these issues may undermine inference about the distribution of zoonotic risk among host taxa (9), and indeed multiple studies have shown that apparent patterns in zoonotic virus richness become insignificant after correcting for total viral richness (13, 14). Yet it remains unclear how this problem could impact more basic scientific questions, including those concerning macroecological patterns in species-level viral diversity.

In this study, we evaluate whether – given the limits of current data – host-level estimates of viral diversity in mammals can be considered biologically meaningful based on their temporal consistency. Even when a species’ total viral diversity has been ground-truthed using a combination of thorough metagenomic sampling and rarefaction-based estimation (15), estimates suggest only ~3-7% of their viruses are captured by existing host-virus association data (10). With such a small proportion of viruses described, it seems plausible that comparative studies of viral diversity are using numbers that are both subject to change and highly sensitive to differences in sampling strategies between different taxonomic groups of hosts and viruses.

We test this hypothesis by analysing a dataset of 6,571 mammal host-virus associations and their year of discovery (defined as the earliest year that a virus was reported in association with a given host), which represents a comprehensive inventory of known associations from 1930 to 2018 (Supp. Figure 1). Our analyses focus on wild mammals, because the historical intensity of pathogen discovery effort on domestic species could confound inference of broader trends across mammals (Supp. Figure 2). Firstly, we examine virus accumulation curves to test whether current absolute viral richness estimates in well-sampled orders and species are likely to be accurate, applying a test borrowed from research on parasite biodiversity (16, 17): richness estimates can only be taken as “stable” – and thus reflective of values close to the truth – if accumulation curves have passed an inflection point towards an asymptote (18). In contrast, if viral diversity is still accumulating exponentially, current estimates may have little correlation to “true” (unknown) viral richness. Secondly, we evaluate the historical stability of relative viral richness estimates across wild mammal host species, families and orders, by testing the rank correlation between present-day and past estimates at annual timesteps. If the correlation of relative viral richness remains fairly stable over time despite changes in sampling effort, this would suggest that species viromes have been sampled proportionally, and thus current data can (despite being incomplete) still provide meaningful comparative information about macroecological patterns in viral diversity across mammals.

## Methods

### Mammal host-virus association data over time

We accessed mammal host-virus records (1,277 mammal species and 1,756 viruses, of which 1,073 are currently ratified by the International Committee on the Taxonomy of Viruses, ICTV) from a taxonomically-reconciled, dynamic, multi-source database that is the most comprehensive data source for host-virus associations (VIRION; https://github.com/viralemergence/virion). VIRION compiles data from several static data sources (6), the NCBI Genbank database, and the USAID PREDICT project database (8). Here, we define a host-virus association based on broad evidence of infection: either serological, PCR-based, or viral isolation. Some records describe recently-discovered viral strains that are not yet resolved to species level; to ensure these do not inflate viral richness estimates, we only included taxonomically-resolved virus species, defined as either ratified by ICTV (n=1,073) or reconciled to the internal viral taxonomy of the PREDICT project (n=683) (8).

We defined the “discovery year” for each unique host-virus pair (n=6,571), as the earliest year a given virus was reported in a given host, based on either date of publication (for literature-based records), accession (for NCBI Nucleotide and GenBank-based records), or sample collection (for records from the USAID PREDICT database). The full database contains host-virus association data up to and including 2021; however, novel association records become notably sparser after 2018 (Supp. Figure 1), likely due to delays between viral sampling and full reporting, including taxonomic assignment (6). We therefore excluded all post-2018 records, as such lags could bias inference about virus discovery rate trends in recent years. To examine temporal trends in publication effort (a proxy for discovery effort), for each host species we also extracted annual counts of virus-related publications (by searching for host species binomial plus all known synonyms *and* “virus” or “viral”) from the PubMed database using the R package ‘rentrez’ (19).

We visualised cumulative virus discovery curves and publication count trends over time at order-level (Supp. Figures 2–3) and across all wild mammal species (Supp. Figure 4). With the exception of individual species-level models (Supp. Figure 5), all subsequent analyses included wild species only (n= 1,246) and excluded domestic and common laboratory species, as these species have been subject to very different discovery processes (Supp. Figure 2).

### Modelling trends in viral discovery rates at order- and species-level

We modelled temporal trends in viral discovery rates by fitting generalised additive models (GAMs) to annual counts of viruses discovered in a given taxon (1930-2018), and with a nonlinear trend of year fitted using penalised thin-plate regression splines in ‘mgcv’ (20). We fitted viral discovery models at the order-level (including the top 8 best-sampled mammal orders, which have the highest known viral richness: Cetartiodactyla, Rodentia, Carnivora, Primates, Chiroptera, Lagomorpha, Perissodactyla and Eulipotyphla), and at the species-level for the top 50 highest viral richness species in our dataset. Virus discovery counts were modelled as a Poisson process for all orders, with the exception of Chiroptera, Rodentia and Primates, which were modelled using a negative binomial likelihood due to high overdispersion in counts in recent years (Figure 1). If discovery curves have reached an inflection point in any species or taxon, we would expect to observe a consistent downward trend in discovery rates in recent years. To test this, we identified time periods showing strong evidence of either an increasing or declining trend, defined as periods during which the 95% confidence interval of the first derivative of the fitted spline does not overlap zero. We also examined the sensitivity of inferences to more conservative definitions of viral diversity by additionally conducting order-level analyses using either only ICTV-ratified virus species or defining viral diversity at the genus level (Supp. Figure 6).

**Figure 1:**
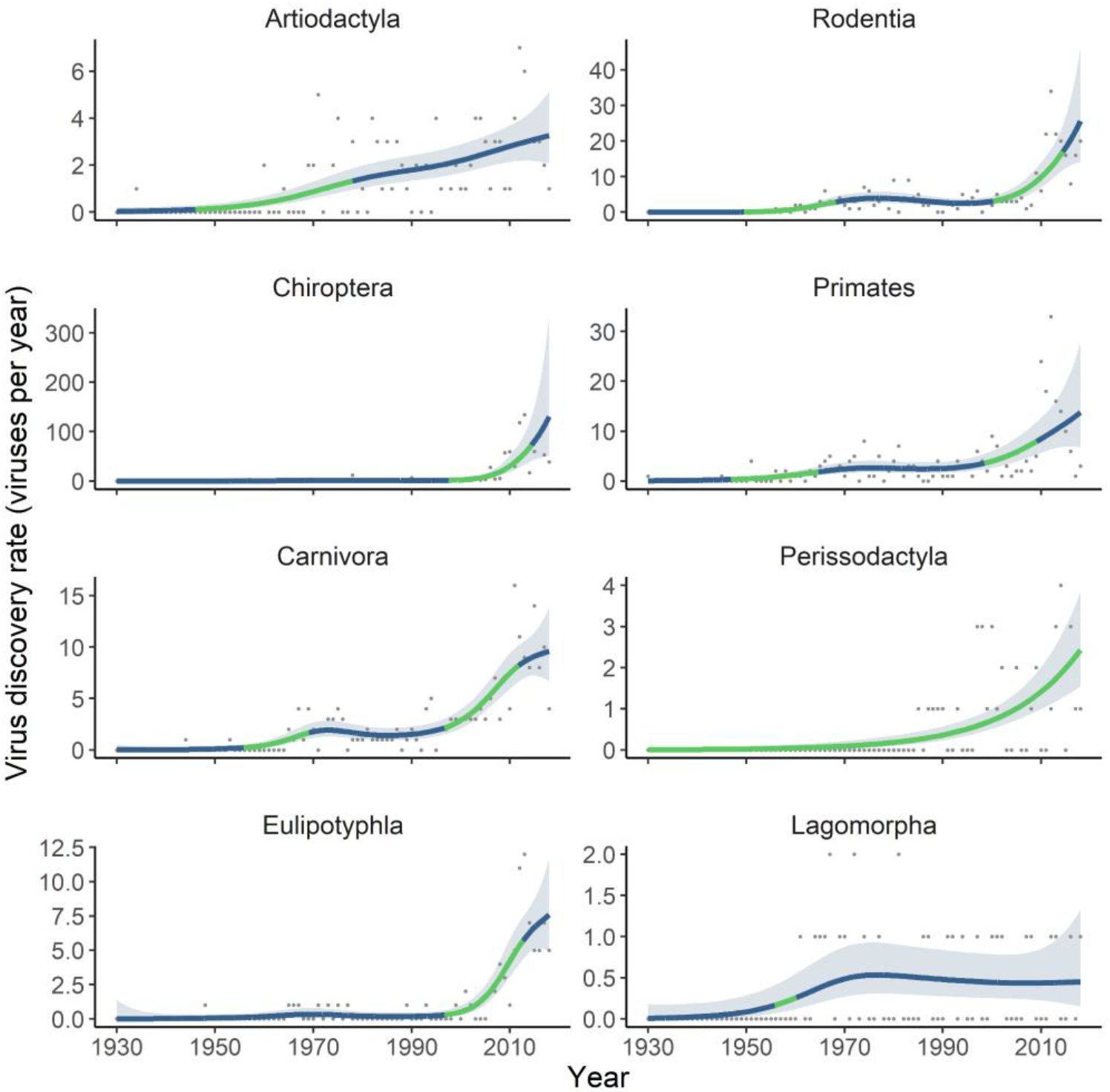
Virus discovery rates within well-sampled mammal orders are still either constant or accelerating. Points show the number of novel viruses discovered per year (1930-2018) infecting wild species of each of the top 8 most virus-diverse mammalian orders. Lines and shading show the fitted temporal trend in virus discovery rate (mean and pointwise 95% confidence interval) estimated using generalised additive models (see Methods). Line colour indicates time periods during which there is either strong evidence of an upward (green) trend in discovery rates (95% confidence interval of the first derivative of the fitted trend not overlapping zero) or no significant trend (blue).

### Evaluating the temporal stability of relative viral richness estimates across taxa

A key untested assumption of most studies that leverage host-virus association data for ecological inference is that, although the mammal virome remains largely uncharacterised, currently known differences in virome composition between species (or higher taxonomic groupings) are nonetheless broadly representative of “true” underlying patterns in viral diversity. If this were the case, estimated differences in relative viral richness across taxa would be expected to stay relatively stable over time, even as discovery effort gradually fills the gaps in species-level virus inventories. Alternatively, unequal distribution of sampling effort across species and time (for example, disproportionate focus on certain host groups of particular zoonotic interest) may severely impact this assumption (9), by causing instability and rapid reordering of viral richness estimates across taxa. We tested this by calculating the rank correlation of viral richness in 2018 to viral richness estimates in annual timesteps backward to 1960 (i.e. comparing the similarity of each annual historical “snapshot” of viral richness to the final snapshot in the study end year) using Spearman’s 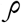. We conducted this analysis at several taxonomic levels, comparing viral richness at the species level (across all mammal species, and separately within each of the key orders listed above), and comparing two different metrics at family and order levels (total viral richness and mean species-level viral richness). We visualised these as curves to examine stability over time. All analyses were conducted in R 4.0.3 (R Core Team, 2020).

## Results

Both cumulative discovery curves and fitted GAMs show that viral discovery in mammals is still in an upward growth phase, with little evidence of discovery rates consistently declining towards zero or viral richness reaching an asymptote for either domestic or wild species (Figure 1; Supp. Figure 2). This trend is mirrored in virus-related publication counts, which are still exponentially increasing year-on-year across most mammal orders and cover an increasingly broad species range over time (Supp. Figure 3). The GAM results show evidence for a general uptick in discovery rates at two main historical junctures (Figure 1). Viral discovery rates first substantially increased during the 1960s, when technological improvements – including density gradient centrifugation for viral isolation, and establishment of the first human diploid fibroblast cell lines and the now-ubiquitous African Green monkey kidney Vero cell line – facilitated industrial-scale production of viruses for research or vaccines (21). Discovery rates again increased sharply throughout the 2000s, which coincided with improvements in molecular techniques for detection and next generation sequencing, and with growing funding for wildlife virus surveillance on zoonotic emergence prevention grounds following the 2002 SARS-CoV epidemic (and a subsequent uptick in research and surveillance effort focused on bats; Supp. Figure 3). The overall picture is the same at the species level, with the estimated mean cumulative viral richness across all wild species still increasing exponentially (Supp. Figure 4) and little evidence of discovery rates declining within even very highly-sampled species (many of which are domestic; Supp. Figure 5). These overall trends are very similar when using several more conservative definitions of viral richness (viral genera, ICTV-ratified viruses, or stricter detection criteria excluding serologic detection; Supp. Figure 6).

A consequence of this accelerating trend in virus discovery is that inference of relative viral richness across species and higher taxonomic levels has been unstable over the last 60 years (Figure 2). Across all mammals, there is a consistent, gradual temporal decay in rank correlation between present-day and historical estimates of total viral richness, with species-level curves declining markedly more steeply than those at higher taxonomic levels (dropping to 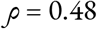 by 1991; Figure 2a). Estimates of mean species-level viral richness at order and family levels (arguably a more relevant metric when considering species contributions to community pathogen maintenance and transmission) are substantially more effort-sensitive than total viral richness, showing much steeper declines (Figure 2a). Within well-sampled mammal orders there is substantial variation in the historical stability of species-level relative viral richness estimates, and results before 1970 become markedly more unstable due to data sparsity in several orders (Figure 2b). Notably, within Chiroptera there has been an extremely rapid reordering of species-level viral richness estimates since 2000 (declining to 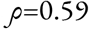 by 2010, and to 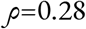 by 2001) as a consequence of the ongoing uptick in research effort (Supp. Figure 3) and viral discovery rates (Figure 1) that occurred after the emergence of SARS-CoV (22).

**Figure 2.**
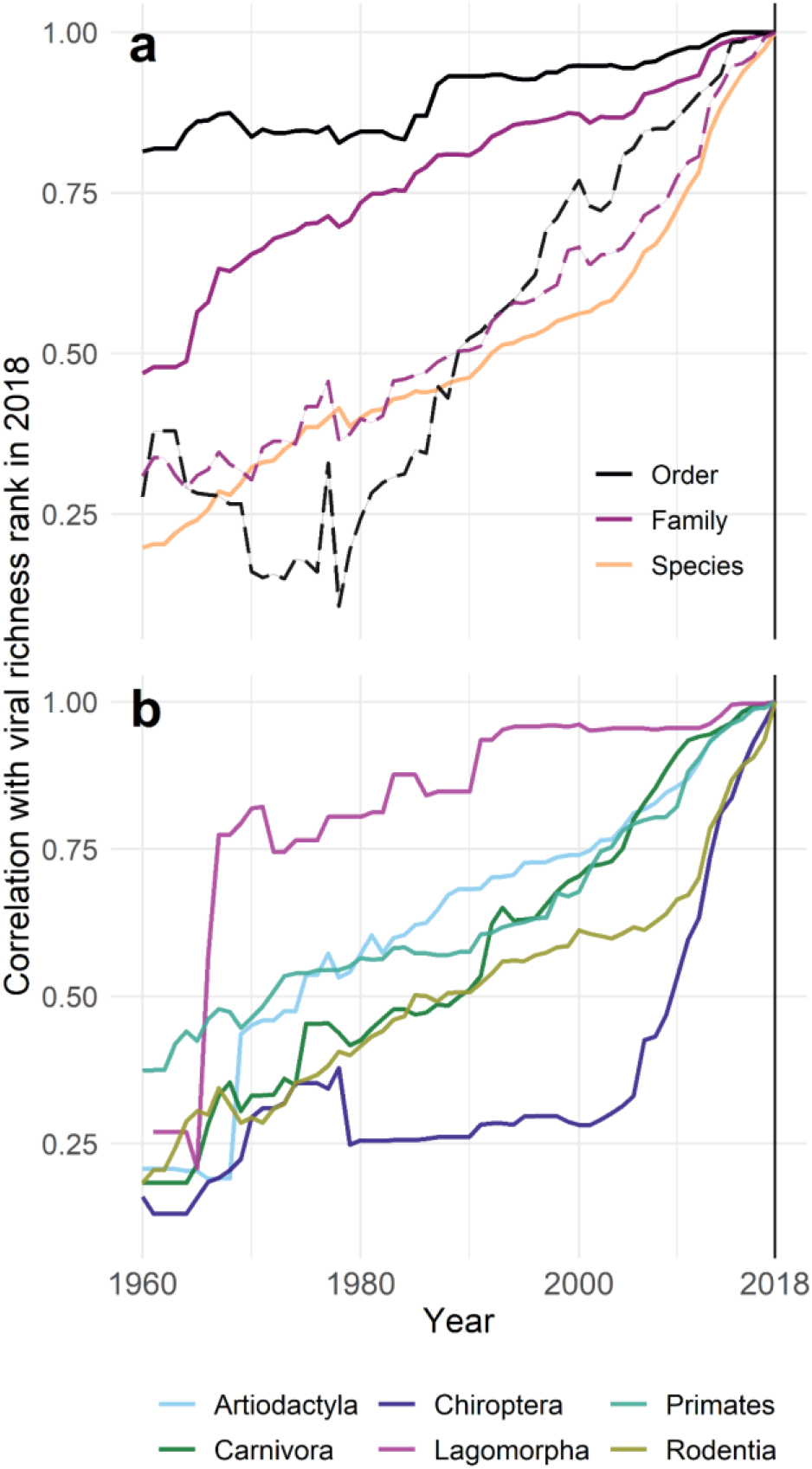
Estimates of relative viral richness across wild mammal taxa are unstable over time. Curves show the rank correlation coefficient (Spearman’s ρ) between viral richness in 2018 (vertical dashed line), and at annual intervals to 1960. Top panel shows curves for all wild mammals (A), comparing viral richness at the species level (n= 1,246), and both total viral richness (solid lines) and mean species-level viral richness (dashed lines) within higher taxonomic groupings (order, n=21; family, n=108) (A). Bottom panel shows separate curves of species-level viral richness within 6 major mammalian orders (B; Artiodactyla, n=153 species; Carnivora, n=148; Chiroptera, n=307; Lagomorpha, n=17; Primates, n=157; Rodentia, n=350). Curve shape denotes temporal stability or instability of relative viral richness estimates across species/taxa; a sharper incline over a given period corresponds to a faster rearrangement of ranked viral richness (i.e. greater instability) in response to discovery effort.

## Discussion

Our results suggest that for the majority of mammal species, viral diversity metrics are still a shifting target, and are largely reflective of historical sampling bias. Given that even the best-studied species do not have fully characterized viromes, these estimates are likely to continue shifting in coming years. Inference made on them, however, might become canonical in the literature – and embed false narratives about viral ecology – if these analyses are not repeated as the global virome becomes better described. The situation might be improved by massively-coordinated, well-funded projects aiming to accelerate viral discovery (11, 23), provided sampling strategies are carefully designed to be taxonomically and geographically representative. However, the rapid recharacterisation of the bat virome that has occurred since the first SARS epidemic highlights a significant risk: if sampling strategies are primarily motivated by either existing (zoonotic) viral diversity estimates or health security concerns linked to specific taxa, such initiatives might only further decouple observed and true underlying viral diversity.

Indeed, the unprecedented general upward trend in wildlife virus discovery effort since 2000 has not been uniformly distributed across taxa and geographical regions. Wild rodents and bats have been particularly heavily sampled and show the highest instability in richness estimates. Ungulates (artiodactyla and perissodactyla) are unique among taxa in that reported viral diversity among domestic species exceeds that detected in wildlife (Supp. Figure 1). While this might reflect the unique ecology of farmed livestock, it is more probable that the data reflect a bias toward sampling from livestock, which poses fewer logistical hurdles than sampling from wild ungulates. Further, many viral discovery efforts focus on detection of targeted viral taxa (e.g. family-level consensus PCR) rather than unbiased approaches that remain cost-prohibitive and analytically challenging. Such evolving detection biases - including renewed efforts to identify bat betacoronaviruses following the emergence of SARS-CoV-2 - could, for example, continue to reinforce the perception of certain host taxa as unusually virus-diverse, although evidence for this remains inconclusive (13). Consequently, it is concerning that a broad comparative study of correlates and geographical patterns of host-virus relationships conducted in 2000 might feasibly have drawn quite different conclusions than a similar study conducted in 2010 or in 2020.

The problem we have identified is not necessarily surprising to many virologists, who have historically been more hesitant to make inference on these limited samples than disease ecologists, and have encouraged particular caution with respect to inference about human health risks (9). Multiple studies have found that correcting for undersampling undermines widespread assumptions about zoonotic risk (13, 14), and we suggest that future studies should similarly attempt to reject the null hypothesis that downstream patterns of zoonotic risk are a neutral consequence of total observed viral diversity. Given that present-day data are a tiny observed subset of the latent “true” host-virus network, there will also likely be value in employing network- or measurement error-based methods that explicitly account for observation biases in analyses of the wildlife virome (24). Overall, since current patterns of host-level viral richness represent an unstable and biased snapshot of the mammal virome, we suggest that inference from host-virus association data needs to be carefully qualified, and may not be a comprehensive foundation for setting future agendas on viral zoonosis research or One Health policy.

## Acknowledgements

The authors were supported by NSF BII 2021909. NM was supported by the Wellcome Trust (217221/Z/19/Z). The authors thank David Redding and the Viral Emergence Research Initiative group for many conversations on the challenges of these data.

## Code and data availability

All the data and code used to generate the results in this article are archived at GitHub (https://github.com/rorygibb/pathogen_discovery/releases/tag/0.1).

## Supplementary Figures

**Supplementary Figure 1:**
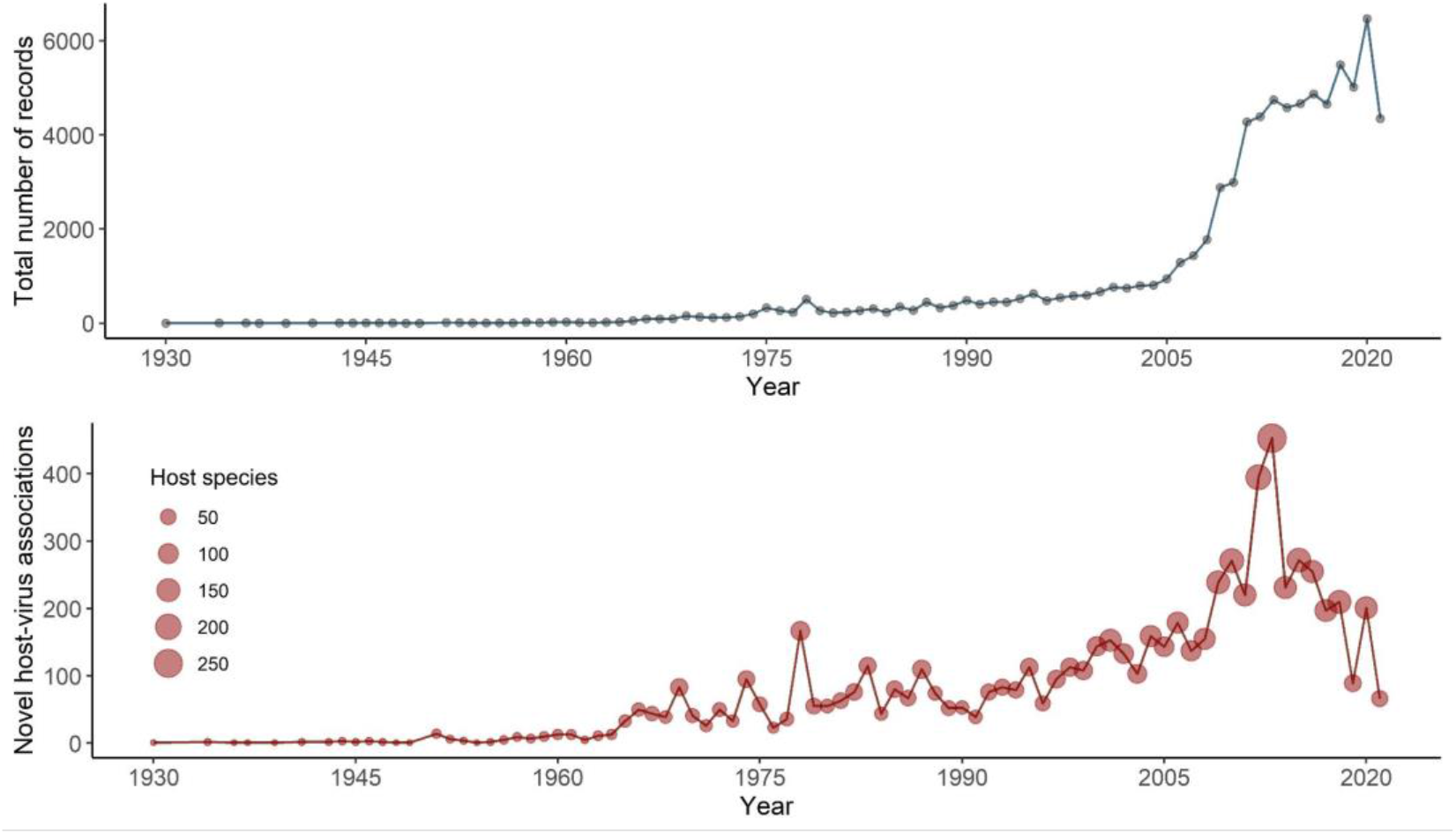
Historical trends in viral discovery in mammals from 1930 to 2021. Top panel shows the annual total number of host-virus association records in the VIRION dataset used for this analysis (see Methods). Bottom panel shows the number of novel host-virus associations reported in each year (i.e. the year in which each unique association was reported for the first time; n=6,571 in total). Point size represents the number of host species in which novel associations were reported per-year. Novel associations show a general decline post-2018 despite the high number of records overall; this discrepancy is likely in part due to lags between detection and reporting (including ICTV ratification; see Methods), so the analyses in this paper only include data up to and including 2018.

**Supplementary Figure 2:**
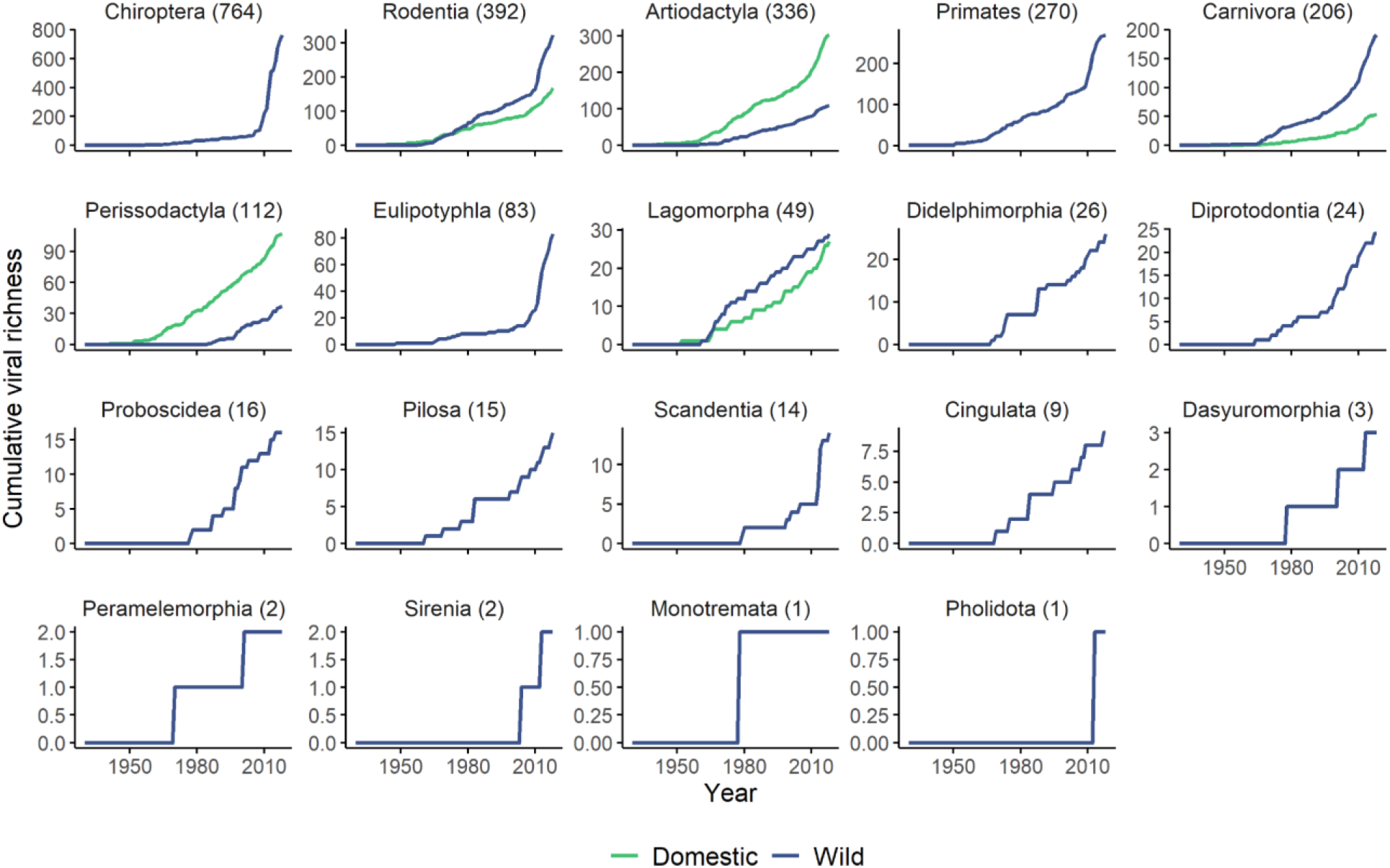
Cumulative viral discovery curves for all mammalian orders linked to at least one virus species in our data. Curves show cumulative viral richness over time across all wild (blue) and domestic (green) species in each mammalian order. Numbers in parentheses denote total known viral richness in each order in 2018, the cut-off date for inclusion in these analyses (Gibb et al. 2021).

**Supplementary Figure 3:**
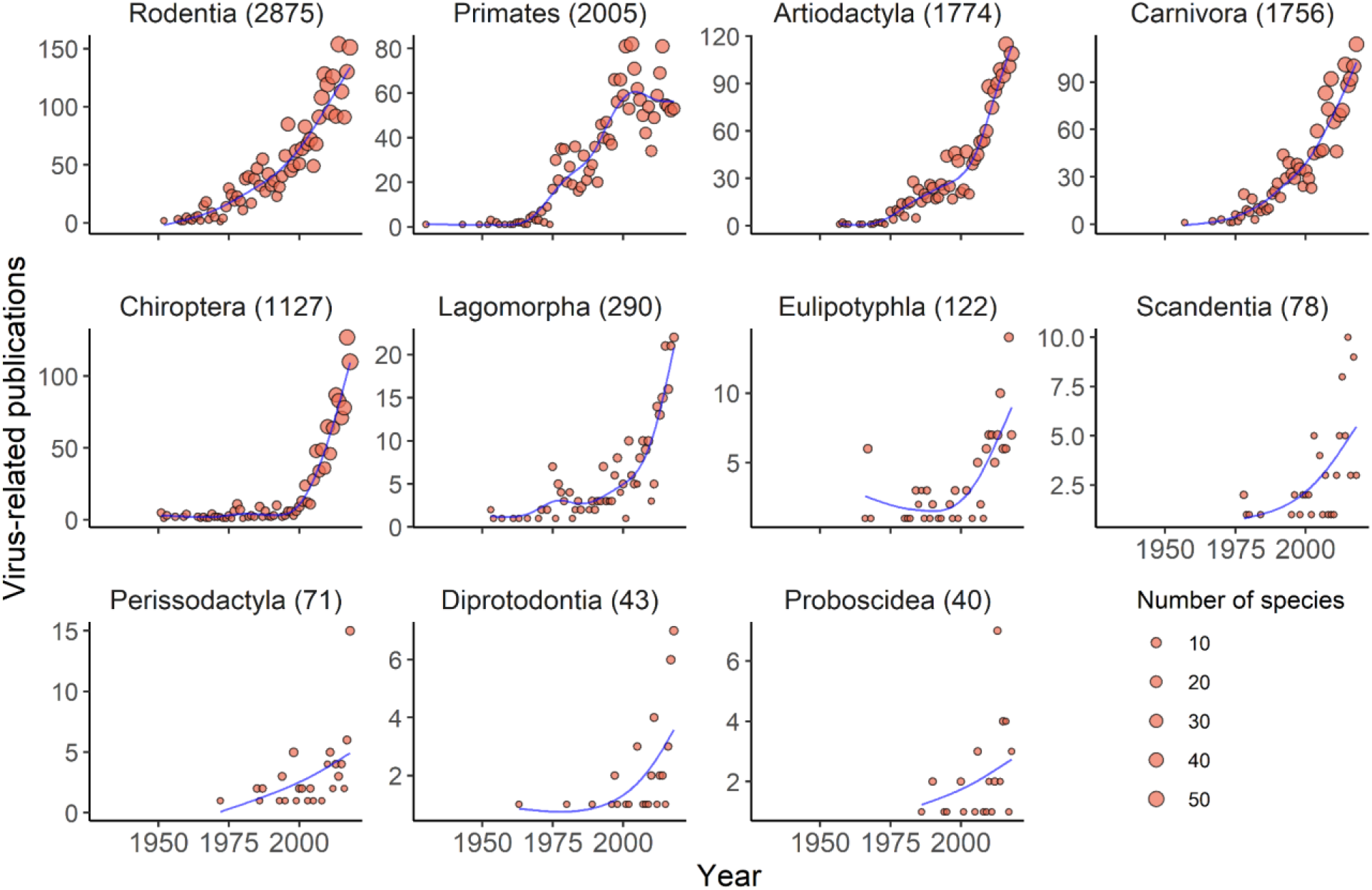
Trends in virus-related publication effort for the most-studied mammalian orders. Points show the annual total number of virus-related publications recorded on PubMed across all wild host species within our dataset (search: “species binomial AND virus OR viral” across the timespan 1930-2018), summed to the order-level. Point size denotes the number of species which had any virus-related publications in that year (i.e. the taxonomic breadth of publication effort). Blue lines are fitted generalised additive model trends, shown for visualisation purposes. Numbers in parentheses denote the total virus-related publications per order in 2018.

**Supplementary Figure 4:**
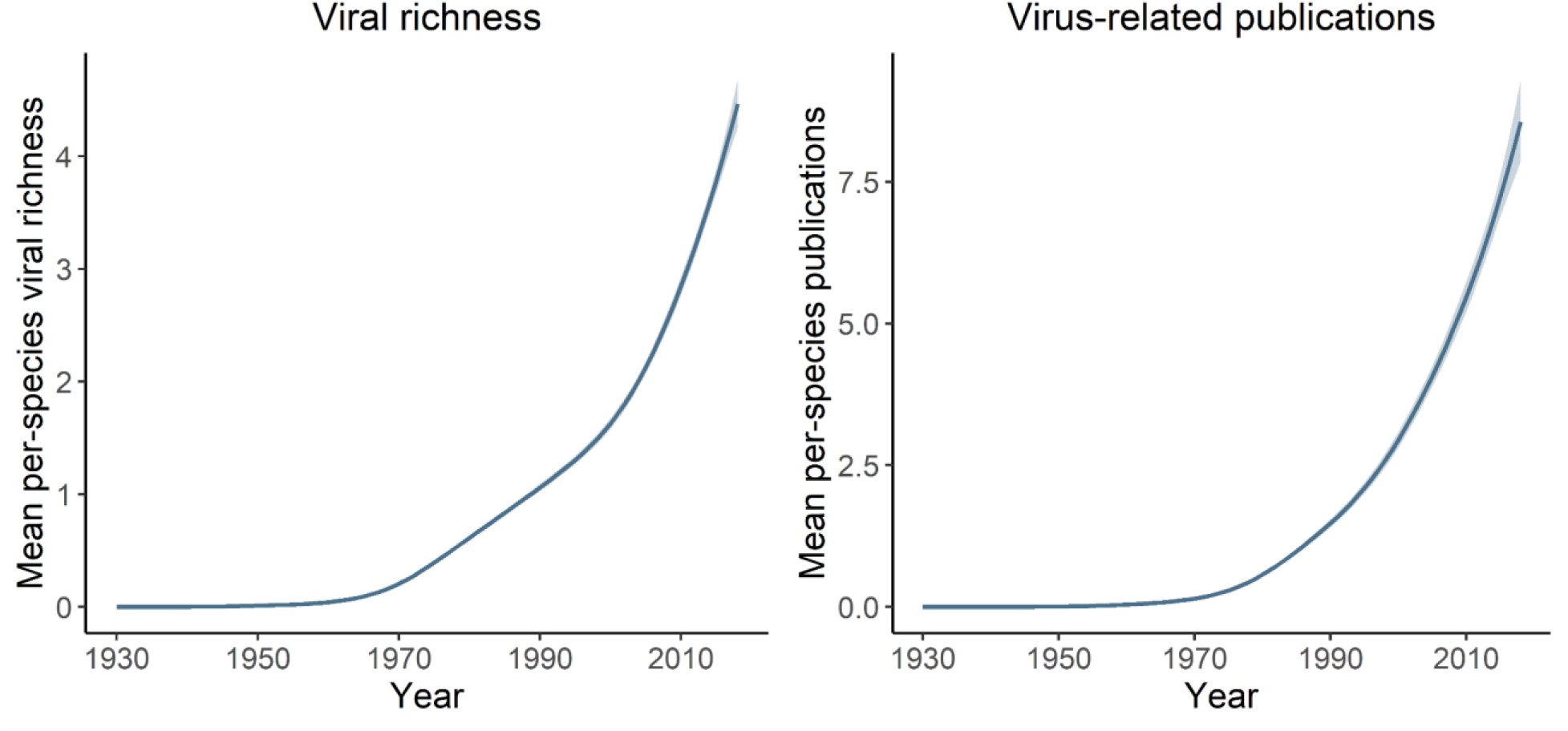
Overall trend in cumulative viral richness and virus-related publication effort at the species-level. Lines and shaded ribbons show the average trend in cumulative species-level viral richness, and virus-related publications from PubMed from 1930 to 2018 (mean and pointwise 95% confidence interval), across all wild mammal species in our dataset (n=1,246). Trends were fitted to annual cumulative counts across all species using generalised additive models with a negative binomial error distribution (to account for overdispersion in counts across species) using ‘mgcv’ (see Methods).

**Supplementary Figure 5:**
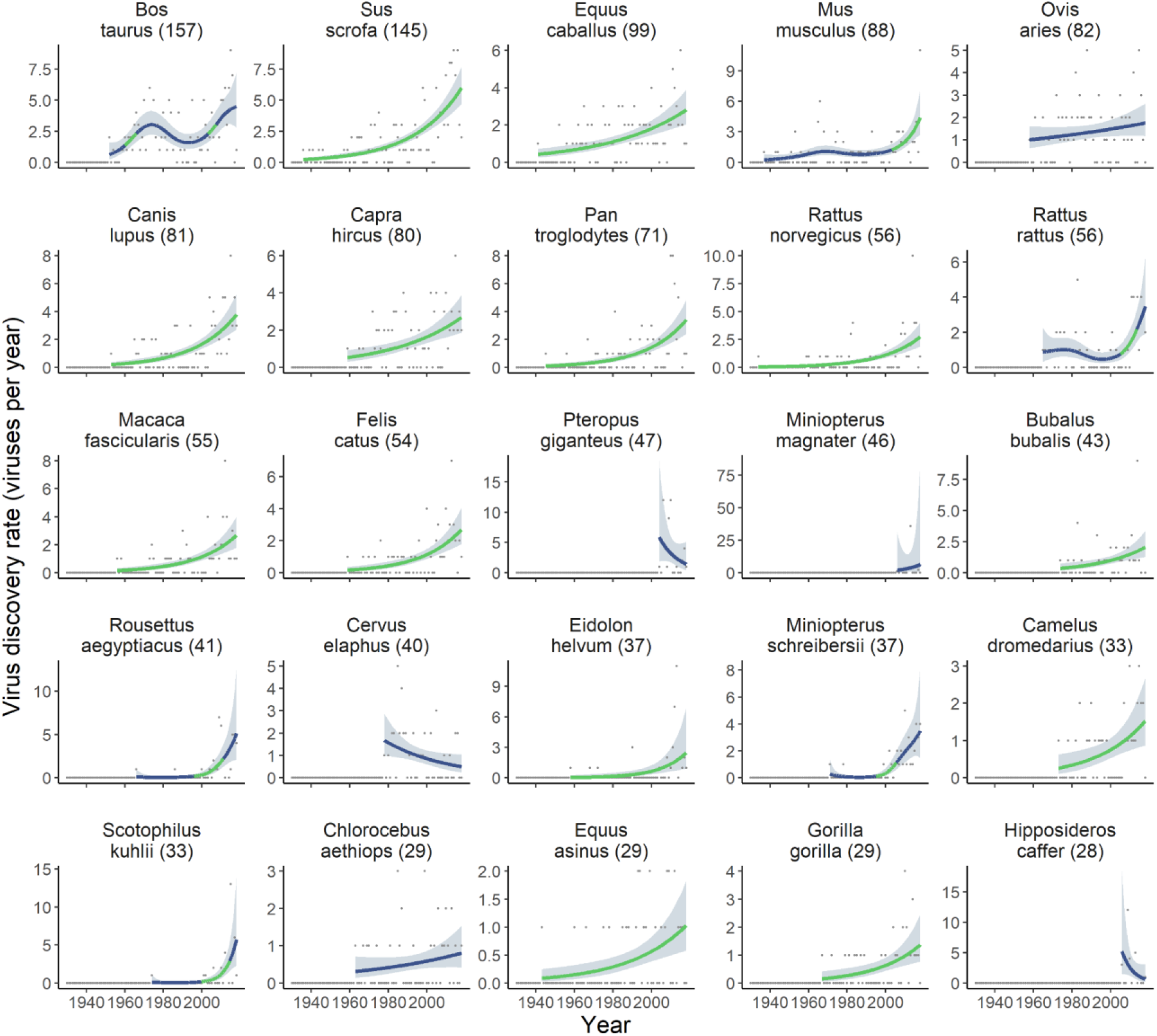
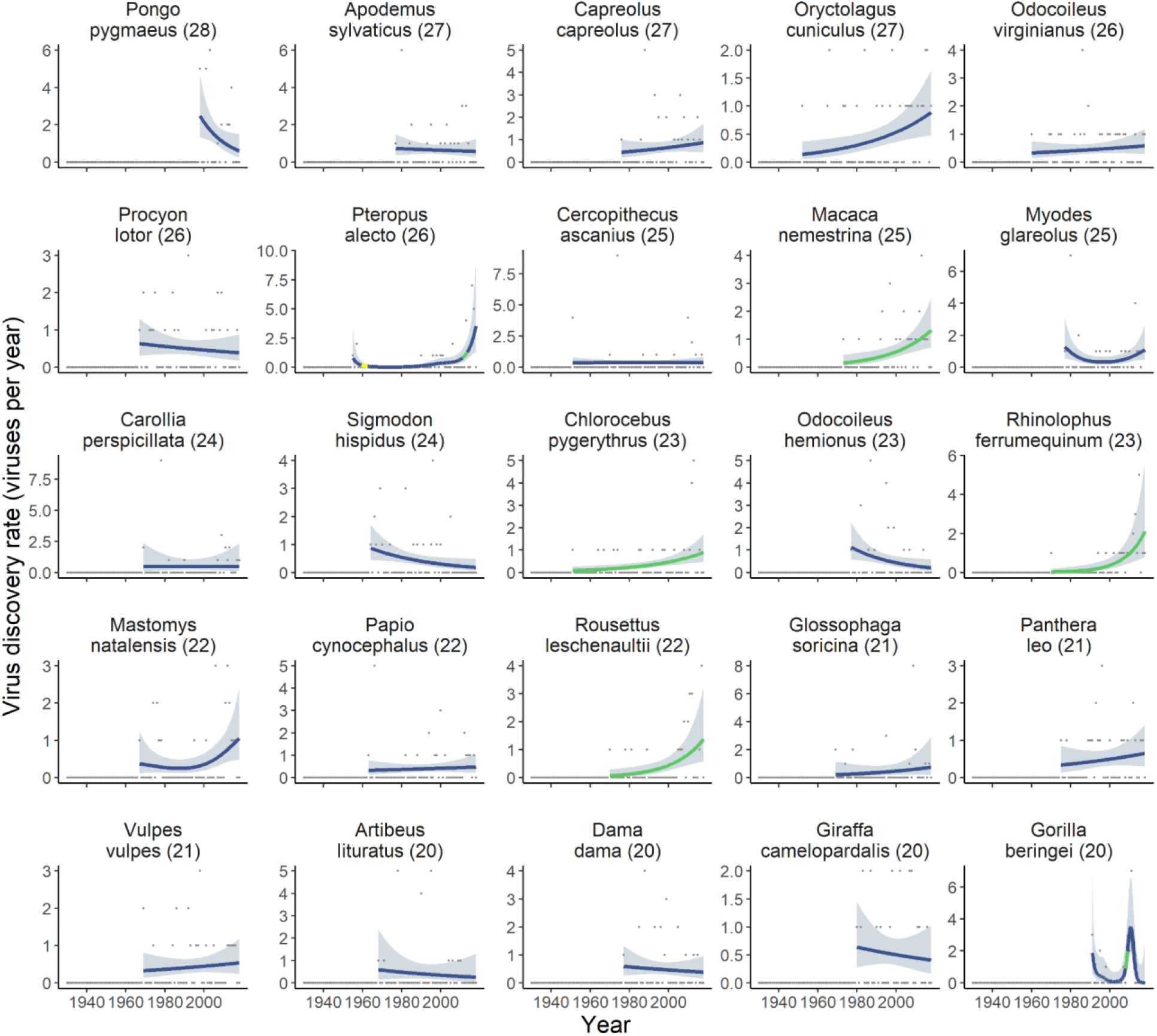
Fitted trends in viral discovery rates for the top 50 highest viral richness mammal species. Points show the number of viruses discovered per year infecting the named mammal species (including both domesticated and wild species), with plot titles showing its total recorded viral richness as of 2018 (in parentheses). Lines and shading show the fitted temporal trend in virus discovery rate (mean and pointwise 95% confidence interval) estimated using Poisson generalised additive models (see Methods). Models were fitted to the time series of annual discovery counts, starting at the year the first virus was reported in a species. Line colour indicates time periods during which there is strong evidence of either an upward (green) or downward (yellow) trend in discovery rates (95% confidence interval of the first derivative of the fitted trend not overlapping zero).

**Supplementary Figure 6:**
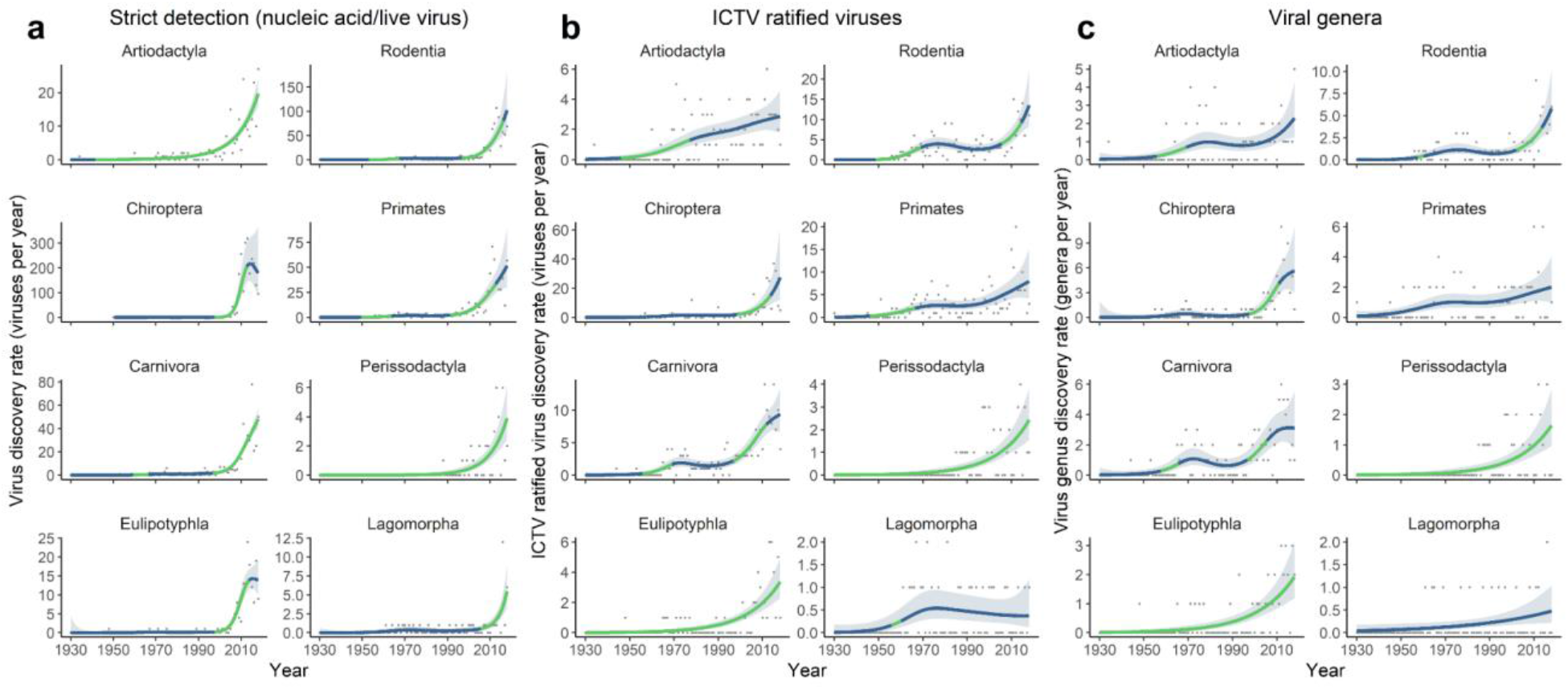
Order-level discovery trends under different taxonomic and detection-based definitions of viral discovery. We repeated discovery curve analyses at mammal order level (Figure 1) based on three alternative, more conservative definitions of viral discovery: detection by either nucleic acid detection, live virus detection or isolation (i.e. excluding serologic detections; A), ICTV-ratified virus species only (B) or viral genera (C) (see Methods for details of generalised additive model fitting). The overall shape of discovery trends is very similar to those based on all virus species (Figure 1), although upward trends under stricter detection criteria are generally steeper (A) and rates in recent years for ICTV-ratified viruses are lower (B), likely due to delays between virus identification and taxonomic ratification (see Introduction).

## Notes

### Competing Interest Statement

The authors have declared no competing interest.

